# Development of a Serum-Free Producer Cell Line Generation Process for Scalable and Efficient rAAV Production for gene therapy applications

**DOI:** 10.1101/2025.10.02.679972

**Authors:** Jose Miguel Escandell, Mafalda Dias, Maria Gonçalves, Daniel Pais, Andreia Conceição, Saurabh Sen, Bruno Figueroa, Karen A. Vincent, Patricia Alves

**Affiliations:** iBET, Instituto de Biologia Experimental e Tecnológica, Apartado 12, 2781-901 Oeiras, Portugal; Instituto de Tecnologia Química e Biológica António Xavier, Universidade Nova de Lisboa, Av. da República, 2780-157 Oeiras, Portugal; Cell Line Development, Mammalian Platform CMC, Sanofi, Framingham, MA USA; Sanofi US Genomic Medicine Unit 225 Second Ave Waltham, MA 02451

**Keywords:** rAAV, Serum free process, rAAV production, Producer Cell lines, Cell line development

## Abstract

The growing field of recombinant Adeno-Associated Virus (rAAV) gene therapy demands scalable and efficient production platforms. Manufacturing platforms based on producer cell lines (PCL) hold substantial promise due to their capacity for high productivity and scalability. However, the generation of stable cell lines for rAAV production is typically performed using serum-containing processes, which pose a risk due to safety and regulatory concerns related to animal-derived components. In this study, we established a fully serum-free process for PCL generation for rAAV production, optimizing critical stages of this process, including transfection methodologies for stable plasmid integration, isolation of highly productive cell pools, and media supplementation to support single-cell cloning.

The developed serum-free process generated clonally derived PCLs with productivity and product quality on par with traditional serum-based methods. This approach significantly reduced the cell line generation timeline and eliminated the challenge related to serum supplementation, adapting it to industry requirements. The reported advances overcome a critical step in cell line development for rAAV production, facilitating the implementation of scalable and efficient production platforms to support the development of rAAV-based gene therapies.

## 2. Introduction

Significant progress has been made in recent years in the development of gene therapy using recombinant adeno-associated viruses (rAAVs). Adeno-associated virus (AAV) belongs to the Parvoviridae family and is characterized by a single-stranded DNA genome of approximately 4.7 kb in size, packaged within a proteinaceous capsid. Notably, these capsids exhibit sequence heterogeneity, each conferring distinct tissue tropisms that enable effective targeting of specific tissues (Zincarelli et al., 2008). AAVs have two genes; r*ep* and *cap* that encode the nine proteins involved in replication, assembly and virus encapsidation (Naso et al., 2017). In rAAV vectors. the viral DNA sequences between the inverted terminal repeats (ITR) have been substituted by a 4.7 kb size DNA expressing the gene of interest. rAAVs are highly versatile and have several advantages: effective gene delivery, a good safety profile, a transient but robust transgene expression *in vivo*, and low immune responses when compared to other vectors (Wang et al., 2024). Therefore, it is not surprising that rAAVs are one of the most commonly used viral vectors in *in vivo* gene therapy and with the greatest potential. By 2025, over 350 clinical trials using recombinant rAAV-based systems had been conducted, leading to the approval of seven gene therapy treatments and several additional treatments are anticipated to emerge in the coming years (reviewed in O. W. Merten, 2024). However, scaling up rAAV production presents significant challenges; Chemistry, Manufacturing, and Control (CMC) processes commonly rely on a transient transfection platform in which HEK293 cells are transfected with a combination of two or three DNA plasmids. A three plasmid transfection platform typically includes a *cis*-acting plasmid encoding the transgene flanked by ITRs, and two *trans*-acting plasmids: one providing the helper functions required for viral replication and packaging, and the other encoding the AAV Rep and Cap proteins(Destro et al., 2024). This method is suitable for small-scale production but presents a challenge for commercial application due to scalability issues and the high cost of raw materials. This bottleneck is particularly pronounced for diseases requiring systemic administration of rAAV, such as muscular dystrophies, where rAAV doses can reach up to 10^15^ vector genomes (vg) per patient (reviewed in Escandell et al 2022). At current volumetric productivities, it is estimated that a 20 L bioreactor would be required per patient, which is not feasible for commercial purposes(A Reid et al., 2024).

Stable producer cell lines (PCLs) offer several advantages compared to other platforms. PCL’s can be fully characterized, they provide higher reproducibility, scalability and efficiency thus streamlining manufacturing workflows, are cost-effective with enhanced genetic stability, and can simplify regulatory approval processes for rAAV products once established. These advantages have the potential to significantly reduce production costs, thereby making rAAV-based gene therapies more accessible and affordable. However, although PCLs offer a valuable alternative for gene therapy manufacturing, these production systems have faced challenges in reaching the commercial stage. The primary limitation lies in the lack of robust cell line development methods necessary to generate PCLs; only a few reports have described protocols for PCL generation tailored to rAAV production (J. Escandell et al., 2023; Martin et al., 2013). Notably, most of these protocols include steps requiring the use of serum-containing media (*e.g.* transfection, single-cell cloning). The presence of serum presents a significant challenge for regulatory compliance due to an increased risk of introducing contaminants, such as adventitious agents.

In this manuscript, we present a novel serum-free PCL generation protocol for rAAV production. Key parameters were optimized to develop PCLs with high productivity, including the transfection method, the selection process, and the composition of the single-cell cloning medium. Using this approach, we generated PCLs producing an rAAV serotype 5 vector with volumetric productivities above 10^11^ vg/mL and with the expected product quality. We believe that the implementation of these innovations represents a step forward in gene therapy manufacturing, holding the potential to expand access to gene therapy treatments for a broader patient population.

## 3. Material and methods

### 3.1 Cells and cell culture

The parental host cells used in this study were previously described as suitable for rAAV production (Chadeuf et al., 2000; J. Escandell et al., 2023; Martin et al., 2013; Tatalick et al., 2005). Parental cells and PCLs were cultured in EX-CELL medium (Sigma-Aldrich, catalog 14591C) supplemented with 6 mM glutamine (Thermofisher, catalog 25030). The supplements evaluated in this study (Supplement I, II, and III) were obtained from Advance Instruments under the catalog numbers RS-1205, RS-1105, and RS-1305, respectively. Cultures were maintained at 37 °C in suspension shake flasks at 125 rpm, under conditions of 5% CO₂ and 80% humidity. Stable transfection of the parental host cells was performed using a plasmid encoding the AAV *rep2/cap5* genes, a vector genome containing either SEAP or green fluorescent protein (GFP) transgene, and a puromycin resistance gene. Producer cells were passaged every 3–4 days while being maintained in culture, for rAAV production, cells were seeded in production vessels after being cultured for three days.

### 3.2 Transfection Efficiency Analysis by Flow Cytometry

Cells were transfected via lipid-mediated transfection with either lipofectamine (ThermoFisher, catalog 18324020) or lipofectamine 3000 (ThermoFisher, catalog L3000001), electroporation (ThermoFisher) and nucleofection (Lonza Bioscience) (Figure 1A) following the manufacturer’s instructions. Forty-eight hours post-transfection, GFP-positive cells were quantified using a BD FACSCelesta flow cytometer, with a minimum of 10,000 events recorded per sample. Data analysis was performed using FlowJo software to determine the percentage of GFP-expressing cells as a measure of transfection efficiency.

**Figure 1.**
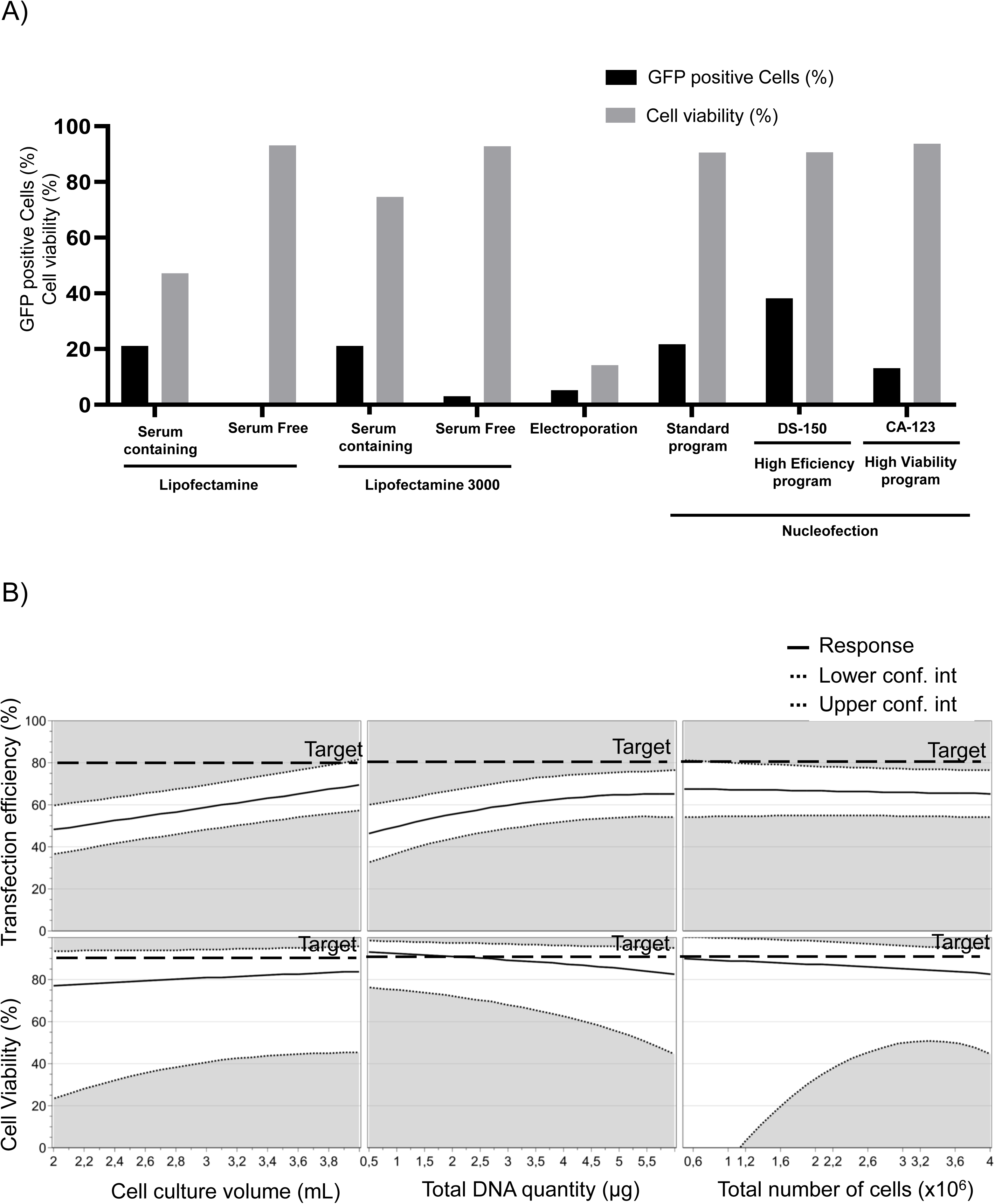
Transfection optimization in serum free chemically defined conditions. (A) Various transfection methods were tested on the parental cell line using a plasmid carrying GFP as transgene. After two days, cells were analyzed by flow cytometry to evaluate transfection efficiency. Cell viability was assessed using trypan blue exclusion method. (B) The optimal transfection method (Nucleofection protocol DS-150) was further optimized using a Design of Experiments (DoE) approach. The prediction plot generated by the DoE model illustrates the relationship between transfection efficiency and cell viability across the tested design space. The black dashed lines (lighter) represent the upper and lower bounds of the model-predicted response with an interval of confidence of 99%, while the black dashed lines (bolder) indicate the predefined target thresholds of 80% transfection efficiency and 90% cell viability. The solid black line shows the predicted trend across varying experimental conditions.

### 3.3 Plasmid Linearization

Plasmid linearization was performed using the restriction enzyme ApaLI or NotI depending on the plasmid (New England Biolabs, catalog R0507S and R3189). Briefly, 10 µg of circular plasmid DNA was incubated with 20 units of restriction enzyme in 1× NEBuffer *Cutsmart* (New England Biolabs, catalog B6004S) at 37 °C for 1 hour. Complete digestion was confirmed by agarose gel electrophoresis. Linearized plasmid was purified and precipitated using ethanol. The DNA pellet was resuspended in 70% ethanol, vortexed briefly, and centrifuged at 12,000 × g for 10 minutes. The ethanol was removed, and the DNA was air-dried under sterile conditions. DNA was resuspended in sterile nuclease-free water and quantified using a Lunatic spectrophotometer (Unchained labs).

### 3.4 rAAV titer determination

Cell lysis was performed using a solution containing 0.5% sodium deoxycholate (Sigma-Aldrich, catalog 302-95-4), 2 mM magnesium chloride (Sigma-Aldrich, catalog 302-95-4), and 60 units of Benzonase (Merck, catalog 101695001), followed by incubation at 37 °C for 90 minutes. To quantify viral genomes, the cell lysates were treated with 60 U/mL DNase (Promega, catalog PROMM6101) for 30 minutes at 37 °C, followed by 2.5 mg/mL Proteinase K (Merck, catalog 3115836001) digestion for an additional 30 minutes at 37 °C. Proteinase K was inactivated by heating at 95 °C for 20 minutes. Quantitative PCR (qPCR) was carried out using a LightCycler 480 (Roche) with SYBR Green detection reagent (Roche, catalog 04707516001). The primers targeting the BGH polyA sequence (Fw: 5′- TCTAGTTGCCAGCCATCTGTTGT-3’; Rv: 5′-TGGGAGTGGCACCTTCCA-3′) were used for rAAV5 quantification.

### 3.5 Design of Experiments for Transfection and media composition optimization

A DOE approach was employed to optimize transfection conditions using MODDE software (Sartorius). Factors investigated included DNA amount, number of cells to be transfected and cell density after transfection. A full factorial design was constructed to evaluate the main effects and interactions between these variables. Response variables included transfection efficiency (percentage of GFP-positive cells) and cell viability.

To optimize media composition for limiting dilution cloning, a DOE was used to design a response surface experiment that investigated the effects of multiple factors, including Supplement concentration (Supplement II, 0–30%), Conditioned Medium (CM) (0–50%), and EX-CELL medium (0–50%) composition. In this case JMP software (SAS Institute) was used to perform the design and model optimization. A base medium of 50% DMEM/F12 supplemented with 6 mM glutamine was maintained in all conditions. This design included 16 runs to assess the interactions and individual effects of the variables. The primary response variables were cell survival and cell outgrowth on day 7 post-seeding, quantified using image-based analysis. For CM preparation, parental cells were seeded into 125 mL shake flasks at an initial density of 0.5×10^6^ cells/mL. Cultures were maintained in an orbital shaker at 125 rpm with a 25 mm orbital diameter at 37 °C and 5% CO₂. After 36–48 hours, the cell suspension was harvested, and cell viability (>95%) and density (1.5–2×10^6^ viable cells/mL) were confirmed. To produce CM, the harvested cell suspension was centrifuged at 300 × g for 5 minutes to remove cells and the supernatant was filtered through a 0.2 μm filter unit. The resulting CM was stored at 4°C for up of two weeks, and used to support single-cell outgrowth in subsequent experiments.

### 3.6 *rep* expression measurement

Cells were transfected by nucleofection and samples were collected at different time points. Total RNA was extracted from cells using the RNeasy Mini Kit (Qiagen, catalog 74104) according to the manufacturer’s instructions. RNA concentrations were assessed using a Lunatic spectrophotometer (Unchained labs). cDNA was synthesized from 2 µg of total RNA using the Transcriptor HiFi cDNA Synthesis Kit (Roche, catalog 5081955001) following the manufacturer’s protocol. The reverse transcription reaction was performed with oligo(dT) primers to ensure high fidelity in cDNA synthesis. r*ep* gene (Forward primer 5’-GGCAGCCTTGATTTGGGA -3’; Reverse primer 5’-GACCAGGCCTCATACATCTCCTT - 3’) expression levels were quantified using qPCR with *HPRT* (Forward primer 5’-CCTGGCGTCGTGATTAGTGAT-3’; Reverse primer 5’-AGACGTTCAGTCCTGTCCATAA-3’) as the housekeeping gene for normalization. The qPCR reaction was performed in a total volume of 20 µL containing 2 µL of cDNA, 10 µL of SYBR Green Master Mix (Roche, catalog 04887352001), 0.5 µM of forward and reverse primers, and nuclease-free water. *rep* expression levels were also normalized by transfection efficiency, assessed in parallel using flow cytometry analysis of GFP expression (as mentioned above).

### 3.7 Full to empty ratio determination

Producer cell cultures were harvested three days post-infection with wild-type adenovirus 5. The cells were lysed using a Tween-20-based lysis buffer as previously described (J. Escandell et al., 2023). The lysate was clarified by centrifugation at 5000 × g for 15 minutes at 4 °C, followed by filtration through a 0.45 μm filter to remove cell debris. The clarified lysate was subjected to purification using PhyTip® columns (PhyNexus) according to the manufacturer’s instructions. The ratio of full to empty vector particles was quantified using mass photometry with a OneMP instrument (Refeyn).

### 3.8 Determination of copy number and sites of integration

To measure plasmid copy number of different PCLs generated with either circular or linear plasmid, genomic DNA was extracted from cell pellets using the High Monarch Genomic DNA Purification Kit (New England BioLabs, catalog T3050L) according to the manufacturer’s protocol. Droplet digital PCR (ddPCR) was used to determine the plasmid copy number. Specific primer and probe sets were designed to target the plasmid of interest and a reference single-copy gene for normalization. BGH poly A sequence Forward primer (5′-TCTAGTTGCCAGCCATCTGTTGT-3’), BGH poly A Reverse primer (5′- TGGGAGTGGCACCTTCCA-3′), BGH poly A Probe (5’-6FAM-TCCCCGTGCCTTCCTTGACC-MGBNFO-3’); E6 Forward Primer (5’ TTCACAACATAGCTGGCACTATAG - 3’), E6 Reverse Primer (5’ - CTGTCGTGCTCGGTTGC - 3’), E6 Probe (5’-6FAM-CCAGTGCCATTCGT-MGBNFO-3’). The ddPCR reaction was prepared in a final volume of 20 µL, containing 10 µL of 2×ddPCR Supermix (Bio-Rad, catalog 1863024), 5 µL of extracted DNA, 900 nM forward and reverse primers, 250 nM fluorescently labeled probe, 5.0 IU restriction enzyme Hind III (New England Biolabs, catalog R3104) and Nuclease-free water to complete the volume. Droplets were generated using the QX200 Droplet Generator (Bio-Rad) and transferred into a PCR plate. The reaction was subjected to the following cycling conditions in a thermal cycler: Initial Denaturation: 95 °C for 10 minutes. Amplification Cycles (×40): Denaturation: 94 °C for 30 seconds. Annealing/Extension: 55 °C for 1 minute. Final Stabilization: 98 °C for 10 minutes. Hold: 4 °C indefinitely. After amplification, droplets were analyzed on the QX200 Droplet Reader (Bio-Rad). Data were processed using QuantaSoft software (Bio-Rad). Copy numbers were normalized to the single-copy reference gene (Albumin) and expressed as copies per genome equivalent. For integration site determination DNA libraries were prepared using the Oxford Nanopore Technologies Ligation Sequencing Kit. Briefly, end-repair and dA-tailing reactions were performed to generate suitable DNA fragments for adapter ligation. Adapter ligation was conducted to enable compatibility with nanopore sequencing flow cells. The prepared libraries were quantified and quality-checked prior to loading. Libraries were loaded onto an R10.4.1 flow cell and sequenced on the Nanopore P2 Solo platform for up to 48 hours. Basecalling was performed live using Dorado integrated into MinKNOW. Plasmid insertion site identification was conducted through chimeric read analysis using the Galaxy platform. Validation was performed by visual inspection of the breakpoint alignments using IGV (Integrative Genomics Viewer).

## 4. Results

PCL generation involves several steps, including transfection, stable plasmid integration, selection for drug resistance, and single-cell isolation to establish a homogeneous cell population. These steps are essential to enhance PCL productivity, platform robustness, reproducibility, and consistency with product quality. While two methods for PCL generation for rAAV in serum-containing systems have been reported using this host (J. Escandell et al., 2023; Martin et al., 2013), the current study aimed to develop a serum-free process for PCL generation to better align with industry standards. As a critical first step, we focused on optimizing plasmid transfection using a serum-free process (Figure 1A and Supplementary figure 1A). A plasmid encoding the *rep2* and *cap2* genes for rAAV production and Green Fluorescent Protein (GFP) as a transgene was used as a model system. Cells were cultured in serum-free medium, and transfection efficiency and cell viability were assessed 48 hours post-transfection. To evaluate transfection performance, three approaches were tested: (1) Lipofection, using Lipofectamine or Lipofectamine 3000, and (2) Electroporation, employing standard electroporation buffers or (3) the Nucleofection platform. While Lipofectamine-based methods worked well under serum-containing conditions, achieving transfection efficiencies of 21% with cell viabilities of 47% (Lipofectamine) and 75% (Lipofectamine 3000), these reagents failed to transfect cells in serum-free conditions, as indicated by the absence of GFP expression after 48 hours. Similarly, Electroporation was ineffective with a low transfection efficiency of 5% and cell viabilities of only 14%. In contrast, Nucleofection demonstrated both high transfection efficiency and high cell viability across the three available programs. The program DS-150 exhibited the highest transfection efficiency (38%) while maintaining optimal cell viability (90%) when compared with the other nucleofection programs (Figure 1A).

To refine Nucleofection parameters to efficiently transfect the cells, we employed a Design of Experiments (DoE) approach to systematically evaluate the effects of 3 variables—total cell number, plasmid DNA quantity, and post-transfection cell culture volume—on transfection efficiency and cell viability. A total of 16 experimental conditions were tested, with transfection efficiency assessed by flow cytometry and cell viability measured using trypan blue exclusion method (Figure 1B).

Optimization of the Nucleofection parameters significantly improved transfection efficiency while maintaining high cell viability. This optimization doubled transfection efficiency from the original non-optimized conditions to over 70%, with cell viability remaining above 90%. The DoE-generated statistical model further validated the optimal transfection conditions, predicting a transfection efficiency of 71.5% (95% CI: 64.5–78.6%) and a cell viability of 88.6% (95% CI: 55.2–97.6%). The observed strong correlation between predicted and measured transfection efficiencies (R=0.98) underscores the robustness and predictive accuracy of the model within the tested parameter space (Supplementary figure 2).

Moreover, the DoE study showed that balancing DNA quantity and cell number increased transfection efficiency; experiments incorporating 6 µg of plasmid DNA and 4 million cells consistently outperformed other conditions, achieving a maximum observed transfection efficiency of 68.8% with a corresponding cell viability of 85.8%. In contrast, conditions with lower DNA quantities (0.5 µg) and smaller cell numbers (0.5 million) resulted in significantly lower transfection efficiencies (as low as 30%) despite maintaining high cell viabilities (>87%).

We next compared the transfection efficiency of circular and linearized plasmid DNA configurations, as plasmid conformation is known to influence transfection outcomes such as integration sites and copy number (Lehner et al., 2013). A plasmid carrying *rep2/cap2* and GFP as the transgene was linearized by ApaLI digestion (Figure 2A) and transfected alongside its circular counterpart (Figure 2B). Different cell populations were transfected using Nucleofection with the optimized protocol previously described. Three days post-transfection, flow cytometry analysis revealed that transfection efficiencies of cells transfected with the circular plasmid DNA were similar to those achieved previously (Figure 1A vs Figure 2B). However, Transfection efficiency was lower when using linearized plasmid DNA compared to its circular counterpart, as shown in Figure 2B (circular: 39% ± 7 SD; linearized: 11% ± 2 SD). while high cell viabilities were maintained in both cases. Although with differing transfection levels, these data suggest that both plasmid configurations— linearized and circular—are suitable for transfecting cells in a serum-free process.

**Figure 2:**
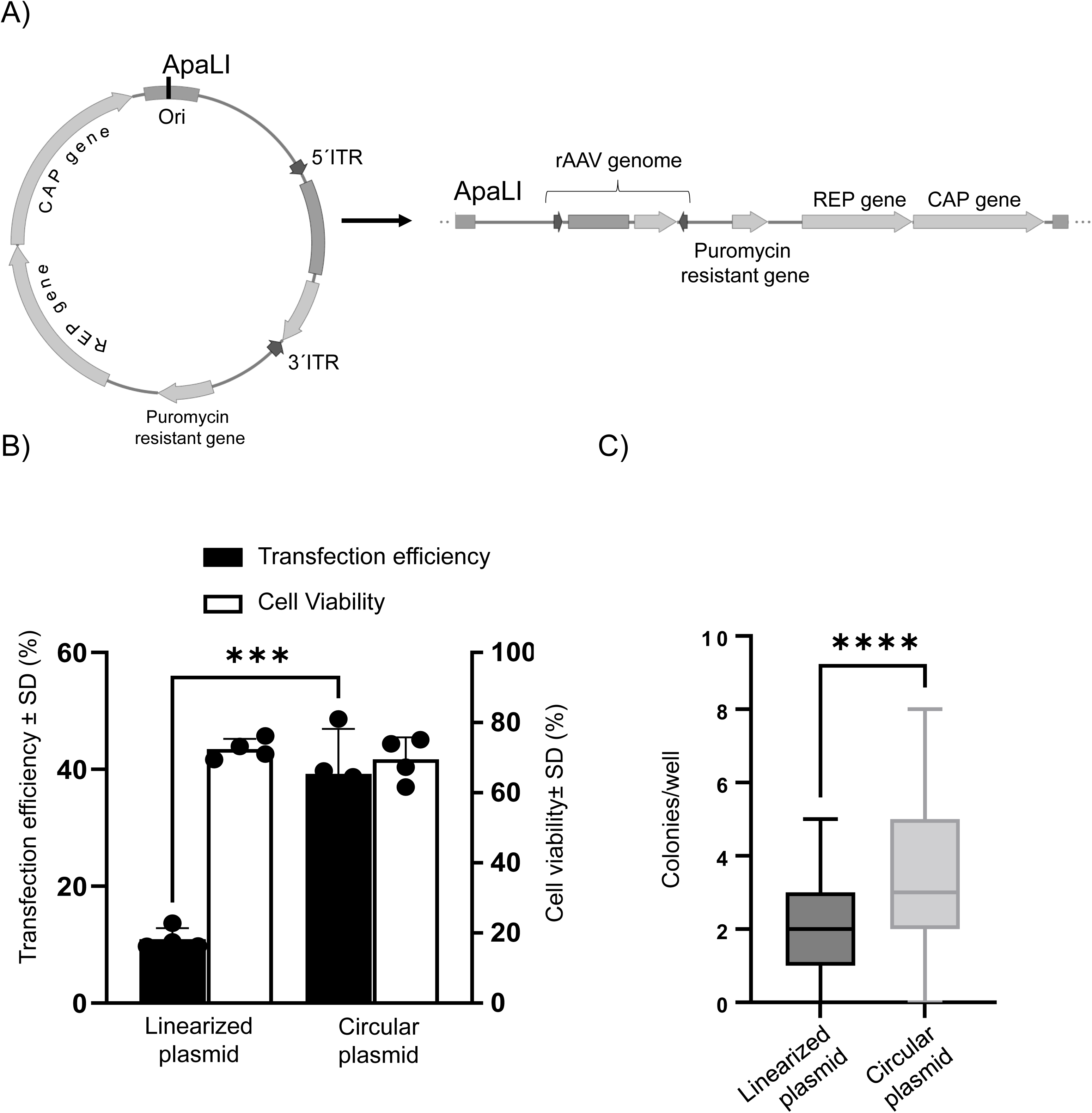
Transfection Comparison of Linear vs. Circular Plasmid Configurations. (A) Schematic representation of a recombinant plasmid to produce rAAV in both circular and linearized forms. The circular plasmid (left) includes key genetic elements: origin of replication (Ori), restriction site used for linearization (ApaLI), antibiotic resistance genes (puromycin resistance), and functional genes (*rep* and *cap*). The linearized form is displayed in the right. (B) Transfection efficiency assessed by flow cytometry three days post-transfection using optimized conditions derived from the DOE in Figure 1. Both circular and linearized plasmids were transfected into parental and clonal cell populations. Cell viability was measured using the Trypan Blue exclusion method. (C) Box-and-whisker plot showing the distribution of viable colonies per well using the Masterwell (MW) approach. Colony formation efficiency following transfection with a plasmid encoding SEAP and rAAV5. Cells were seeded at 4,000 cells/well in 96-well plates using EXCELL Medium with 0.2 µg/mL puromycin. After 15 days, colonies were imaged and counted. The graph shows the minimum, maximum, and median colony numbers per well. Statistical analysis was performed using a non-parametric t-test; Asterisk (***) and (****) indicates statistical significance of (p < 0.001) and (p < 0.0001), respectively.

After optimizing the transfection method in serum-free conditions, we evaluated different selection strategies (Supplementary figure 1B). Selection was initiated three days post-transfection. Two selection methods were tested: the Masterwell (MW) approach, based on described protocols (J. Escandell et al., 2023; Martin et al., 2013) where cells are subjected to selection in a 96-well plate, and a suspension-based approach which involved culturing cells post-transfection in suspension with puromycin at 0.2 µg/mL. While the suspension-based selection offered advantages such as enabling single-cell deposition after selection and faster PCL generation, this protocol failed to generate a stable cell population (data not shown). We hypothesized that the lack of cell adherence and the selective pressure in early suspension culture may have compromised cell viability and recovery, ultimately preventing the expansion of a puromycin-resistant population—an effect likely exacerbated by the physical and metabolic stress associated with selection in suspension conditions, such as those encountered in shake flask cultures. The MW approach, however, consistently generated robust and uniform wells with selected populations. Transfected cells were seeded at 4,000 cells per well in EX-CELL Medium with 0.2 µg/mL puromycin, resulting in the formation of viable colonies ranging from 1 to 10 colonies per well visible at 14 days post seeding. Viable cell outgrowth was achieved using both linear and circular plasmids with this method, further emphasizing the reliability of the MW approach for the generation of stable transfected pools (Figure 2C).

Since both plasmid configurations supported viable cell outgrowth using the MW approach, we next evaluated the impact on the ability to isolate high-producing PCLs. The parental cell line was transfected with a plasmid (either circular or linearized) encoding *rep2/cap5* and containing a rAAV vector genome harboring the secreted alkaline phosphatase (SEAP) transgene. Following puromycin selection, over 150 wells per condition were screened for rAAV5 productivity. Remarkably, MW pools transfected with the circular plasmid demonstrated up to a 7-fold higher productivity compared to those transfected with the linear plasmid. The cells transfected with circular plasmid achieved titers of up to 7 × 10¹⁰ vg/mL with specific productivities exceeding 10⁵ vg/cell (Figure 3A, 3B), suggesting that transfection with a circular plasmid is a better approach to achieve PCLs with high productivity. Figure 3C highlights the MWs generated by both methods that surpassed production thresholds of 10^8^ vg/mL (39 % of the wells transfected with the circular plasmid vs 6 % for the linear plasmid).

**Figure 3:**
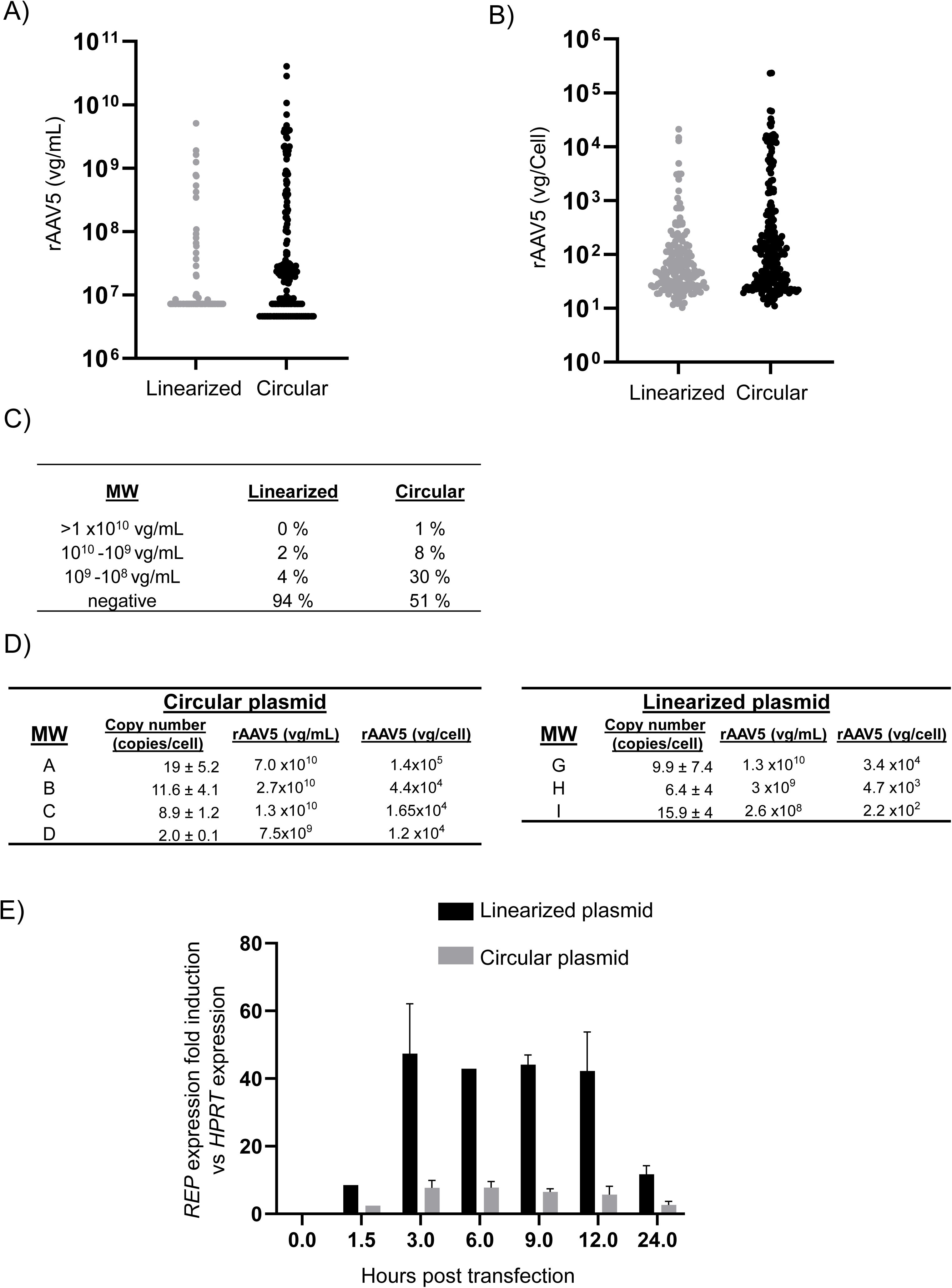
Characterization of Linear vs. Circular Plasmid Configurations in MW Generation. A clone derivative was transfected with rAAV5 carrying the gene encoding the SEAP enzyme via Nucleofection using either linear or circular plasmids. (A) rAAV5 titers are shown as viral genomes per milliliter (vg/mL), and (B) as viral genomes per cell (vg/cell); both assays were performed using production in 96-well format. (C) A summary of rAAV productivity is shown in linearized vs circular plasmid configuration, indicating the distribution of MWs across percentiles. (D) DNA was extracted from the most productive MW populations—generated in shake flask format—and plasmid copy number was assessed by comparing the BGH sequence to the endogenous Albumin gene. (E) Transfected cells were harvested at various time points (0–24 hours), and RNA was extracted. *rep* expression levels were quantified by RT-qPCR, normalized to the housekeeping gene *HPRT*, and adjusted for transfection efficiency. Error bars represent technical variation from two independent cDNA syntheses generated from the same RNA extraction.

We hypothesized that the circular plasmid might exhibit a more favourable integration profile than the linearized plasmid, leading to higher plasmid integration and copy number. To investigate this, plasmid copy number was measured through droplet digital PCR, using primer/probes targeting the BGH polyA sequence in the vector genome, and the albumin gene in the host genome as a control for normalization.

In MWs generated with the circular plasmid, higher rAAV production was generally linked to higher plasmid copy numbers. The most productive MW also had the highest copy number (19 copies per cell; Figure 3D), suggesting a possible linear relationship. This trend was not seen with the linear plasmid. Overall, plasmid copy numbers were similar between both plasmid types, indicating that factors (perhaps site of integration or copy number per site) other than total copy number likely contribute to the higher productivity of the circular plasmid.

Rep protein has been shown to play a critical role in AAVS1 site-specific integration and rAAV production (Howden et al., 2008). To analyse if *rep* gene expression is impaired by linear plasmid transfection, *rep* expression was analyzed via RT-PCR at different timepoints post-transfection in cells transfected with both plasmid configurations (Figure 3E and Supplementary figure 3). Surprisingly, at early time points (3–12 hours), cells transfected with the linear plasmid exhibited significantly higher *rep* gene expression, compared to those transfected with the circular plasmid. These data suggest that, although *rep* expression is crucial for specific integration at the rAAV locus, the lower productivity observed with the linearized plasmid configuration of MW is not due to reduced Rep protein levels post-transfection.

To further assess whether plasmid configuration affects Rep-dependent integration in the AAVS1 locus, we performed integration site analysis using Nanopore sequencing. This approach aimed to determine if differences in site-specific integration could account for the variation in rAAV productivity observed between linear and circular plasmid formats (Supplementary figure 4). Integration events were detected in both conditions (MW A and MW C for circular plasmids and MW I for linear plasmids), indicating that both linear and circular plasmids are capable of mediating site-specific integration in the AAVS1 locus.

These findings suggest that the reduced productivity observed with the linear plasmid is not attributable to a failure in AAVS1 targeting or *rep* expression after transfection.

To meet regulatory requirements, clonality is essential for cell line platforms. Consequently, it was necessary to optimize the process for generation of clones via single-cell deposition in the context of the serum-free generation protocol (Supplementary figure 1C). Initially, single-cell deposition was performed in EX-CELL medium, either alone or mixed with DMEM/F12 and CM, which contains survival-promoting factors secreted by actively growing cells. These factors help support cell survival and proliferation under stressful conditions, such as those encountered during single-cell cloning (U. M. Lim et al., 2013). Despite this, this combination failed to support cell growth (data not shown). Therefore, to promote single-cell growth, various commercially available supplements were evaluated. Supplement II (Figure 4A) was identified as the most effective, supporting a plating efficiency—defined as the percentage of single cells that generated outgrowth—of up to 65%.

**Figure 4.**
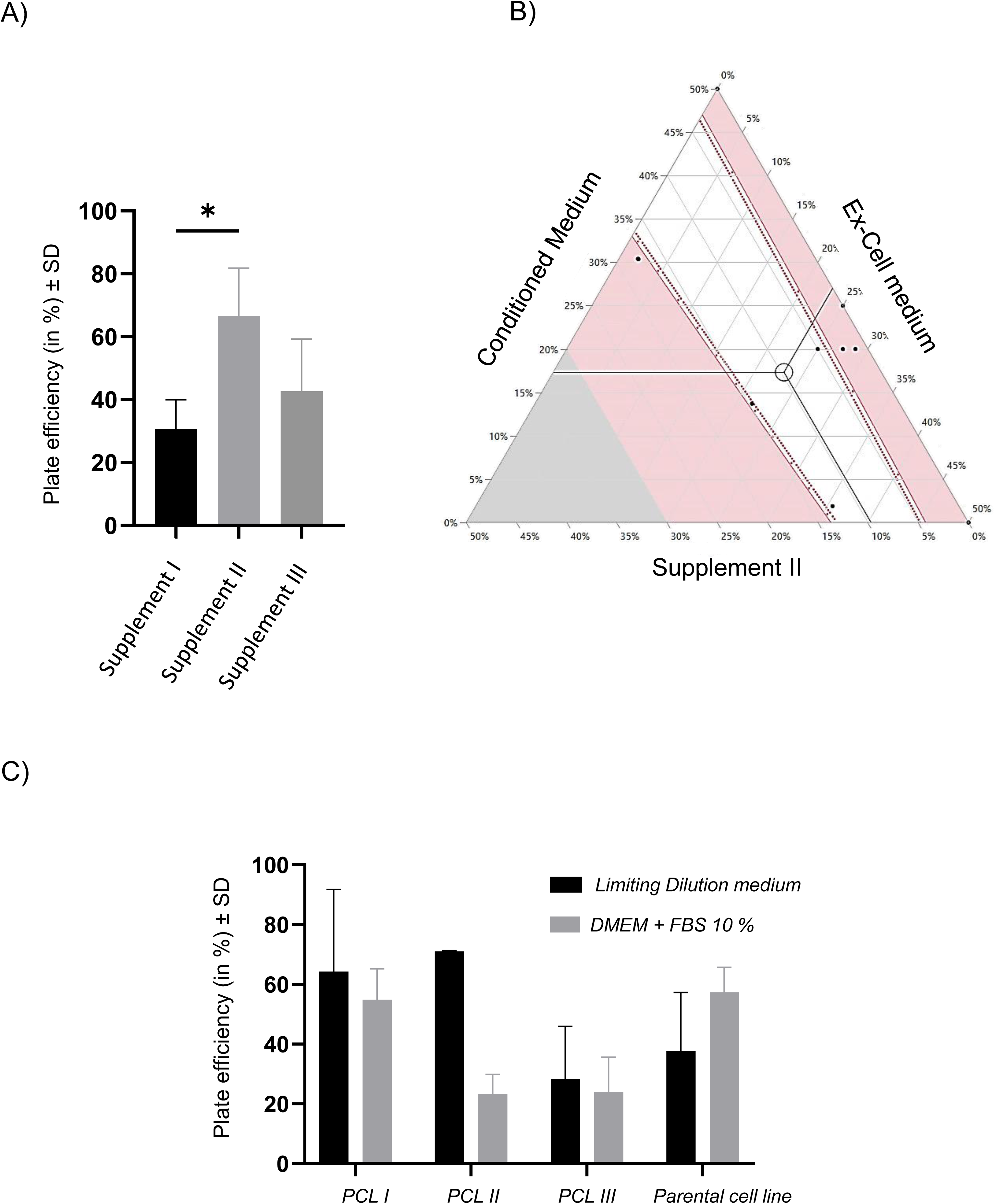
Optimization of Serum-Free Medium Composition for Single-Cell Colony Outgrowth. (A) Comparative analysis of various supplements to determine the optimal supplementation for supporting single-cell colony outgrowth under serum-free conditions. The bar graph depicts the plate efficiency (percentage ± SD) of single-cell colony formation when cultured with Supplement I (black), Supplement II (light gray), or Supplement III (dark gray). Asterisk (*) indicates statistical significance (p < 0.05). Each condition was tested in triplicate across n=3 independent experiments. Plate efficiency was determined by calculating the percentage of single cells that successfully formed colonies after at least 7 days of culture under serum-free conditions. (B) DOE study for medium composition optimization. Three parameters were included in the analysis: percentage of Supplement II, EX-CELL addition, and CM. Different possible mixture combinations and their effect on the plate efficiency are represented, considering values of each component from 0 to 50% (v/v). The remaining 50% medium consists of DMEM/F12. The mixture combinations where the minimum plate efficiency value of 50% on day 7 is achieved are represented by the white area; the red contour (red line and dotted red line) represents the threshold of 50% plate efficiency. The points correspond to the performed experimental mixture combinations. For clarification purposes, in the image it is represented the condition with 10% (v/v) Supplement II, ∼17% (v/v) CM and ∼23% (v/v) EX-CELL medium. (C) Validation of the optimized serum-free condition (limiting dilution medium) in comparison to serum-containing medium for single-cell colony outgrowth. Four different populations (Producer cell lines were labelled as PCLs) were tested, and their outgrowth performance in both media was evaluated.

A DoE approach (Figure 4B) was then used to optimize outgrowth conditions. Fourteen conditions were tested across a range of medium compositions. Colony formation was assessed on day 7 post-seeding. Our model predictions revealed a significant quadratic effect of all medium components on plating efficiency, with a predicted optimal medium composition of 40% CM, 10% Supplement II (2x the manufacturer recommendation), no EX-CELL medium and 50% DMEM/F12 (Supplementary figure 5). As shown in Figure 4B, a ternary mixture profiler illustrates the effect of varying the proportions of CM, EX-CELL medium, and Supplement II on plating efficiency, highlighting in the white area the medium composition mix with plate efficiencies above 50%.

Our experiments confirmed the model’s predictions. Although the highest outgrowth was observed with increased levels of Supplement II, the medium selected for single-cell cloning consisted of 50% DMEM/F12, 25% EX-CELL medium, 20% CM, and 5% Supplement II. This composition was cost-effective, while consistently supporting high plating efficiency, with the model predicting a plating efficiency of 63% and experimental validation showing 60 ± 3%.

Importantly, no residual dependence on Supplement II or CM was observed during subsequent cell expansion (data not shown).

The robustness of this optimized method was validated by comparing cell outgrowth in serum-containing versus serum-free processes. Several producer cell lines (three MW-derived) and the parental host were seeded in both DMEM + 10% FBS and the newly formulated Limiting dilution cloning (LDC) medium (Figure 4C). The serum-free medium composition used for single-cell outgrowth demonstrated comparable plating efficiency across all conditions when compared to FBS-based LDC. In particular, PCL II exhibited a significant increase in seeding efficiency, rising from 20% in FBS to 70% with LDC medium. These results confirm that outgrowth of single-cell clones is feasible with this formulation, facilitating the generation of clonal PCLs in serum free, chemically defined medium conditions.

Supplementary figure 1 outlines the complete protocol for generating PCLs with clonal origin. High-producing MWs transfected with circular plasmid encoding an AAV5-SEAP vector, as shown in Figure 3A, were seeded as single cells. After single-cell deposition and outgrowth, isolated clones derived from the MWs were assessed. These clones were tested for production of the AAV5-SEAP vector, with productivities exceeding 10¹¹ vg/mL (Figure 5A) and specific productivity exceeding 5×10^5^ vg/cell (Figure 5B). Both volumetric productivity and specific productivity were improved after single-cell isolation, likely due to the non-clonal nature of the parental MWs.

**Figure 5.**
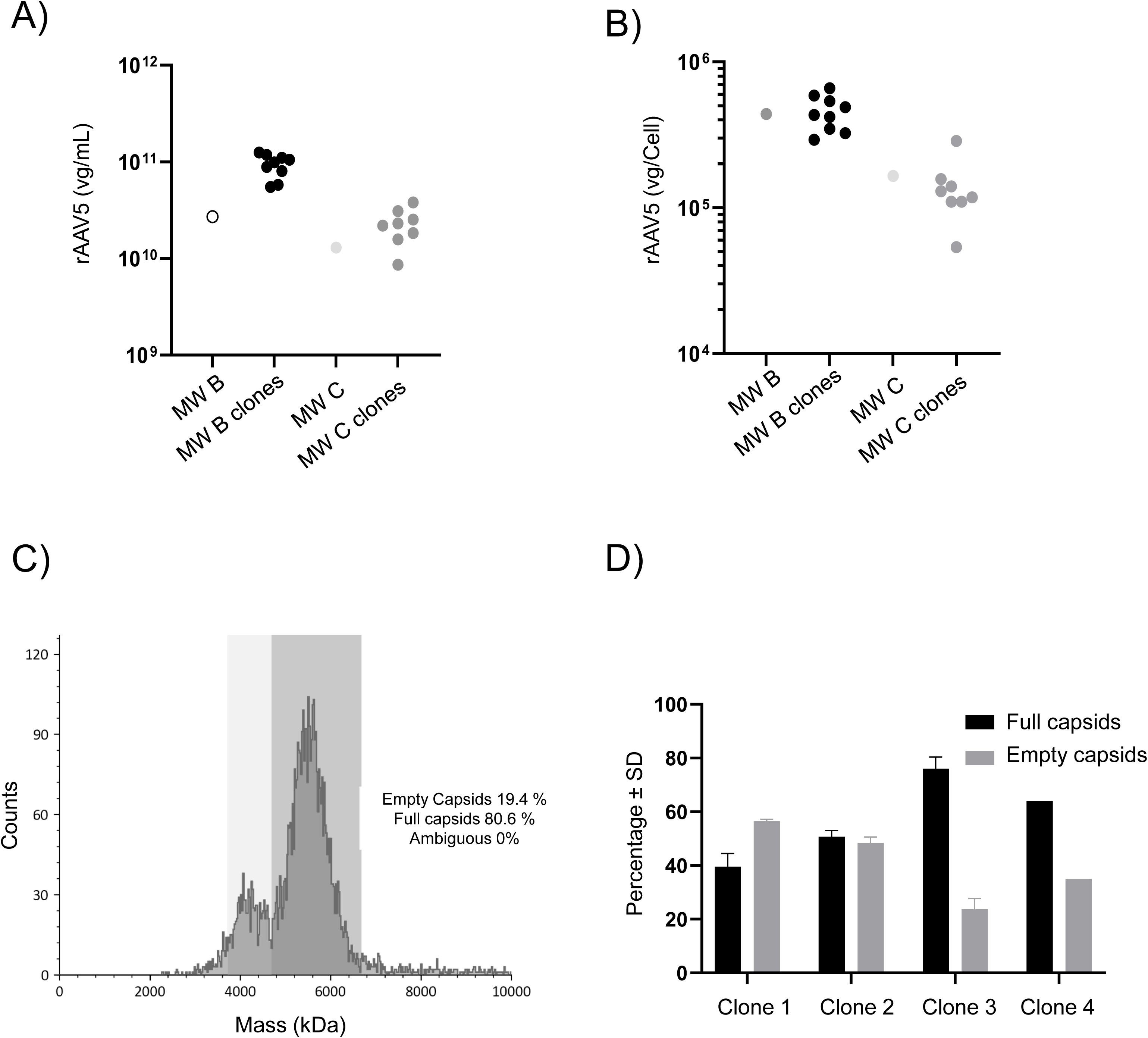
Generation of clonally-derived producer cell line in serum-free conditions. A clone derived from the parental cell line was nucleofected with circular plasmid DNA and subjected to stable transfection under pooled selection conditions. The most productive MW identified in Figure 3A were seeded into limiting dilution cloning medium. The resulting colonies were screened for rAAV5 productivity. The graph compares rAAV volumetric productivity (A) and rAAV vector genome production per cell (B) across the Masterwell stage and Clone stage. (C) The top-producing clones were further analyzed for product quality, focusing on full-to-empty capsid ratio. Mass photometry was used to assess this quality attribute. (D) Variability in the percentage of full and empty capsids among the top 4 clones.

To evaluate the quality of the vectors generated using this serum-free process, rAAV generated from 4 randomly selected clones (from two different MW origins) were purified and analyzed for full-to-empty capsid ratios using mass photometry (Figure 5C and 5D). Different isolated clones produced varying product qualities, with full-to-empty ratios ranging from 40% to 73%. This highlights the importance of screening several clones for product quality determination, as some clones exhibited significantly better full-to-empty ratios than others.

## 5. Discussion

Major bottlenecks in gene therapy manufacturing and consequent application include the lack of scalability and high costs associated with current state-of-the-art methods for viral vector production, reviewed in Escandell et al 2022 (J. M. Escandell et al., 2022). Although PCLs are widely used for manufacturing biologics, such as monoclonal antibodies in Chinese Hamster Ovary (CHO) cell line-based production platforms, their application in rAAV manufacturing is still in its early stages. Of note, none of the eight approved gene therapy products based in rAAV use PCLs as manufacturing platform(Merten, 2024). Therefore, developing a robust and efficient methodology for PCL generation is essential to establish a safe, reliable and cost-effective platform for rAAV manufacturing. In this study, we established a serum-free protocol for generating stable PCLs for rAAV production, addressing critical regulatory barriers inherent to traditional serum-dependent methods. While prior PCL development relied on serum-containing processes (J. Escandell et al., 2023; Martin et al., 2013), such approaches introduce manufacturing risks including lot-to-lot variability of serum and a risk of contamination from adventitious agents (Barone et al., 2020). Using this platform, we consistently generated PCLs with titers exceeding 10¹¹ vg/mL and robust product quality, with a full-to-empty capsid ratio of approximately 70%. Although a different rAAV serotype was used, these titers are comparable to those previously reported for similar PCL platforms (reviewed in Merten, 2024).

A critical step for cell line generation is the transfection of plasmid DNA into the cells. Through a DOE approach, we achieved transfection efficiencies of up to 70% in a serum-free process using nucleofection. This efficiency is comparable to that reported in other studies employing Nucleofection for other challenging-to-transfect cells. For instance, a study using the Nucleofection system achieved a transfection efficiency of 71% in chicken primordial germ cells under feeder-and serum-free conditions (Zhao et al., 2025). Moreover, a study utilizing the Nucleofector technology reported transfection efficiencies of approximately 60% with CHO-K1 cells cultured in serum-free, chemically defined, animal component-free medium (Lang et al., 2016).

During the optimization of the transfection protocol, the use of circular vs linear plasmid DNA was compared and linearization was found to significantly impact rAAV production. Although both formats supported the generation of PCLs, transfection with circular plasmids resulted in 5- to 7-fold higher rAAV titers compared to linear plasmids. Notably, the transfection efficiency of circular plasmids was also higher, resulting in more colonies per well (Figure 2C). These findings contrast with those reported by Lim et al.(S. Lim et al., 2023), who found higher integration rates with linear plasmids. The discrepancies likely arise from differences in plasmid constructs and transfection contexts; in our system, integration of plasmids encoding the AAV genes is an active Rep-dependent process. It was shown that transient expression of the Rep protein following transfection promotes efficient integration and guides plasmid insertion into the AAVS1 locus—a genomic region known to support stable, high-level rAAV production (Luo et al., 2017). The superior performance of circular plasmids may be explained by their suitability as templates for Rep-mediated integration. Henckaerts et al. (Henckaerts et al., 2009) proposed a mechanism in which Rep-mediated site-specific integration involves rolling-circle-like replication and partial duplication of the target locus, a process that may facilitate tandem integration events via template switching—a pathway more accessible with circular templates. Interestingly, we observed that transient Rep expression was higher following transfection with linear plasmids compared to circular ones. This suggests that Rep protein levels are not the limiting factor for integration or productivity. Rather, it reinforces the hypothesis that the structural configuration of the plasmid template— specifically, the circular topology—plays a critical role in enabling efficient Rep-mediated integration and subsequent rAAV production.

Long-read sequencing of high-producing clones confirmed integration at the AAVS1 locus in clones derived from both circular and linear plasmid transfections, indicating that targeted integration can occur regardless of plasmid topology. However, prior studies, such as Tsunoda et al. (2000) (Tsunoda et al., 2000) reported that linearized plasmids are poor substrates for Rep-mediated targeting and integrate less frequently into AAVS1. Our dataset is limited to seven sequenced clones (four from circular and three from linear plasmid transfections). All cell lines selected for analysis were high producers, which may introduce bias toward a more favourable integration profile. In contrast, Tsunoda and colleagues applied a different methodology that enabled analysis of over 80 clones, providing a broader and statistically more robust view of integration behaviour. Nevertheless, in agreement with Tsunoda et al (2000), our results demonstrate that linearized plasmids are able to integrate at the AAVS1 site. Although additional studies with larger clone sets are needed to fully evaluate the influence of plasmid topology on integration efficiency and site specificity after nucleofection, these data support the use of circular plasmids as the preferred format for the stable integration of rAAV components, offering both higher productivity and potentially greater integration efficiency through mechanisms intrinsic to their topology.

Another key step in cell line development is the isolation of single-cell clones. The use of clonal cell lines, as specified by regulatory guidance, significantly enhances product robustness, homogeneity, and quality, thereby improving overall safety(Han et al., 2022). However, outgrowth from a single cell seeded in a well of a tissue culture plate presents challenges and requires optimization. The DOE experiment gave us several insights regarding the optimal single-cell growth conditions in this system. By incorporating CM and supplements, we successfully optimized the process to facilitate cell outgrowth and allowing for the generation of PCLs with high productivity. The optimal medium defined in our study— 20% CM, 5% Supplement II, 25% EX-CELL, and 50% DMEM/F12— supports robust and reproducible results. While the medium preparation is somewhat complex due to the use of CM and supplements, this level of complexity is standard as it reflects the need to address the unique challenges of supporting survival and proliferation of isolated cells (in serum-free conditions)(U. M. Lim et al., 2013). CM provides essential growth factors and cytokines secreted by feeder cells, compensating for the absence of cell-cell interactions in isolated single cells, while defined supplements ensure reproducibility and mitigate batch variability. This dual approach aligns with established methodologies observed in stem cell and primary cell cultures, where tailored media formulations are critical for clonal expansion (Leopold et al., 2019; Singh, 2019). Our findings also indicate that the concentration of Supplement II can be increased to reduce or even eliminate the need for CM. While our study supports the transition away from these variable components that could introduce lot-to-lot variability, cost-benefit considerations remain essential due to the higher expenses associated with elevated concentrations of commercial supplements.

Despite its advantages, the platform still presents certain limitations. At present, the workflow involves a two-step process comprising an initial MW pool selection phase followed by single-cell cloning. To improve efficiency and reduce timelines, future developments will focus on refining the selection strategy to enable direct enrichment of high-producing cells in suspension prior to clonal isolation, similar to CHO cell line development platforms used for monoclonal antibody production (Abali et al., 2024). Nonetheless, even in its current form, the platform outperforms transient systems by offering a more robust, scalable, and cost-effective solution for rAAV manufacturing. Moreover, further integration of automation and artificial intelligence holds promise for streamlining this workflow, enhancing consistency, and addressing the bottlenecks of this platform by enabling data-driven selection and reducing manual intervention (Plante K, 2025).

In summary, this work demonstrates the successful generation of PCLs with clonal origin under serum-free conditions, achieved through the optimization of multiple steps in the cell line development workflow. This approach not only aligns with regulatory expectations for clonality and process consistency but also ensures high productivity and good product quality. Moreover, it enables the development of a scalable and cost-effective rAAV manufacturing platform within a defined and accelerated timeline.

## 6. Data availability

The research data used and methods shown in the main and supplemental figures are available from the lead contact upon reasonable request.

## 7. Acknowledgments

The authors acknowledge the Gulbenkian Institute for Molecular Medicine Genomic Services team for their technical expertise and support in performing nanopore sequencing and related data analyses. Images were generated with Biorender.com.

## 9. Author contributions

J.M.E., writing – original draft, project conceptualization, project supervision, investigation, methodology. M.D., investigation, methodology, visualization, and formal analysis. M.G., investigation, methodology, visualization, and formal analysis. D.P., investigation, methodology, visualization, and formal analysis. A.C., investigation, methodology, visualization, and formal analysis, S.S., validation, and resources, B.F., project supervision and funding acquisition. K.V., project conceptualization, project administration, funding acquisition and supervision. P.G A. project supervision, project conceptualization, project administration, and funding acquisition. All authors: writing – review & editing.

---

**Supplementary Figure 1.**
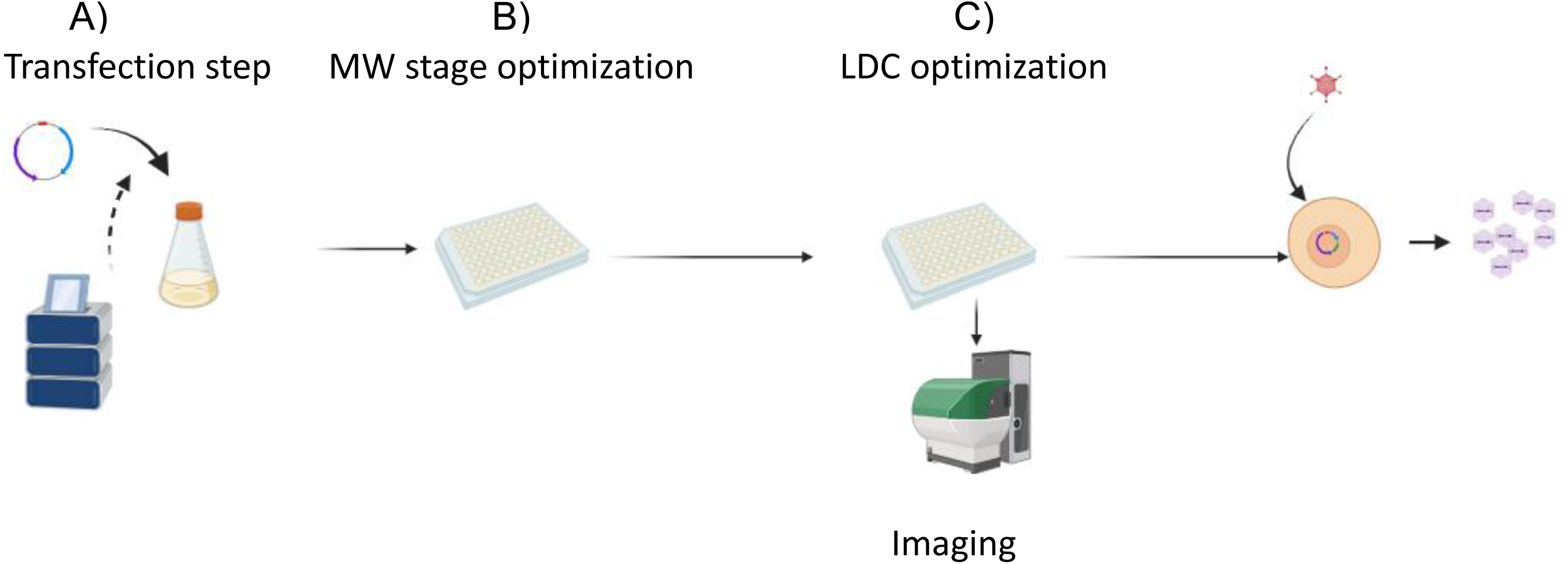
Serum-Free process for producer cell line generation. This manuscript describes the optimization of several parameters critical for establishing a serum-free process for PCL generation. The image illustrates key steps of the process, including: A) Transfection method optimization. B) Selection processes by MW stage method. C) Limiting dilution cloning (LDC) medium optimization to ensure single-cell outgrowth.

**Supplementary Figure 2.**
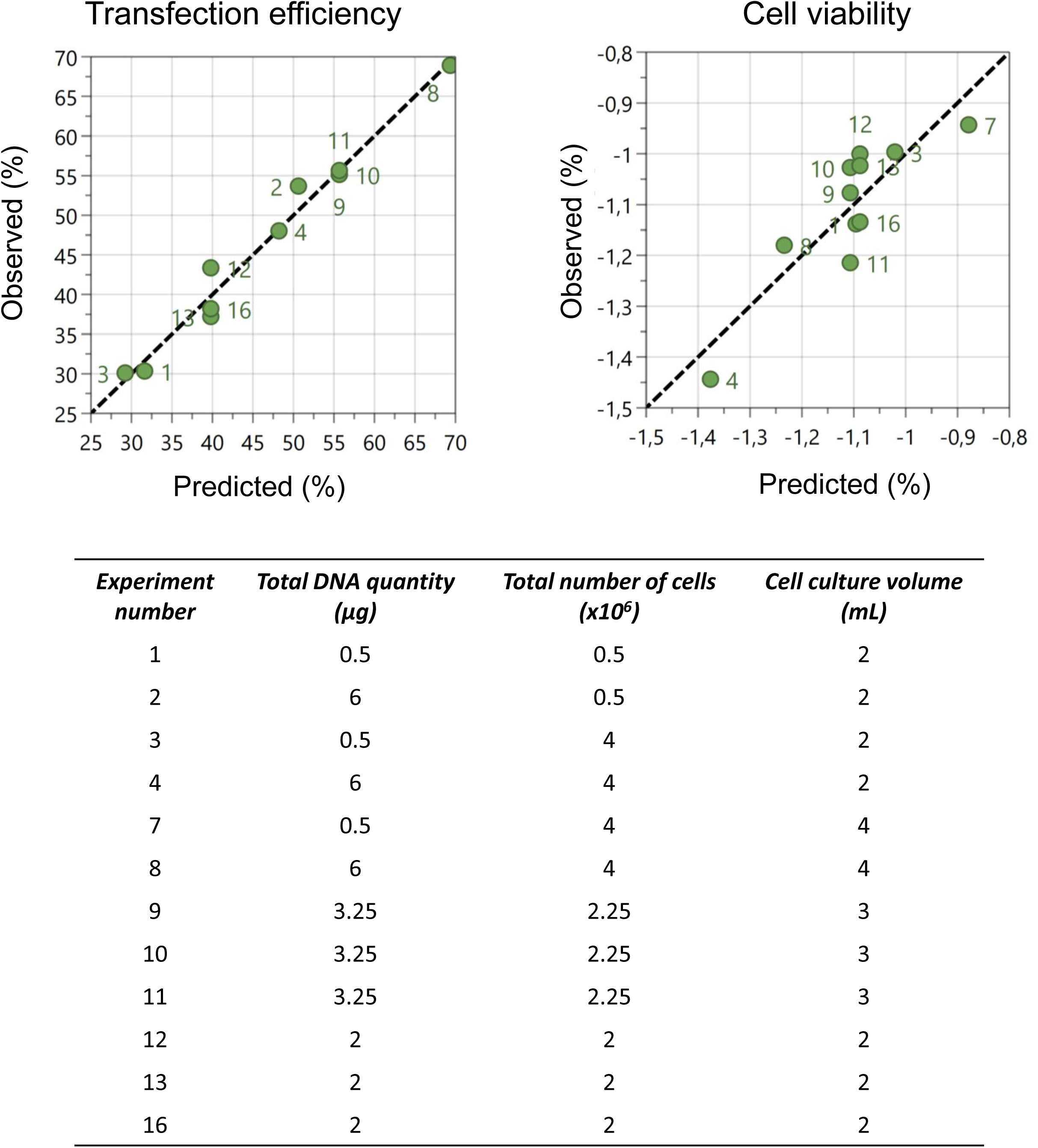
Correlation between transfection efficiency, cell viability, and DOE predictions. The correlation between transfection efficiency and cell viability (empirical data) and values predicted by DOE using MODDE software is shown. Transfection efficiency was measured 48 hours post-transfection by flow cytometry, while cell viability was assessed using the trypan blue exclusion method. The graph displays observed vs. predicted values for both transfection efficiency and cell viability conditions. For the cell viability, a negative log transformation was applied, as suggested by the MODDE software. The green dots on the prediction plots, numbered between 1-16, represent the different experimental conditions. Statistical analysis for transfection efficiency: N = 11, degrees of freedom (DF) = 6, R² = 0.98. For cell viability: N = 11, DF = 5, R² = 0.77.

**Supplementary Figure 3.**
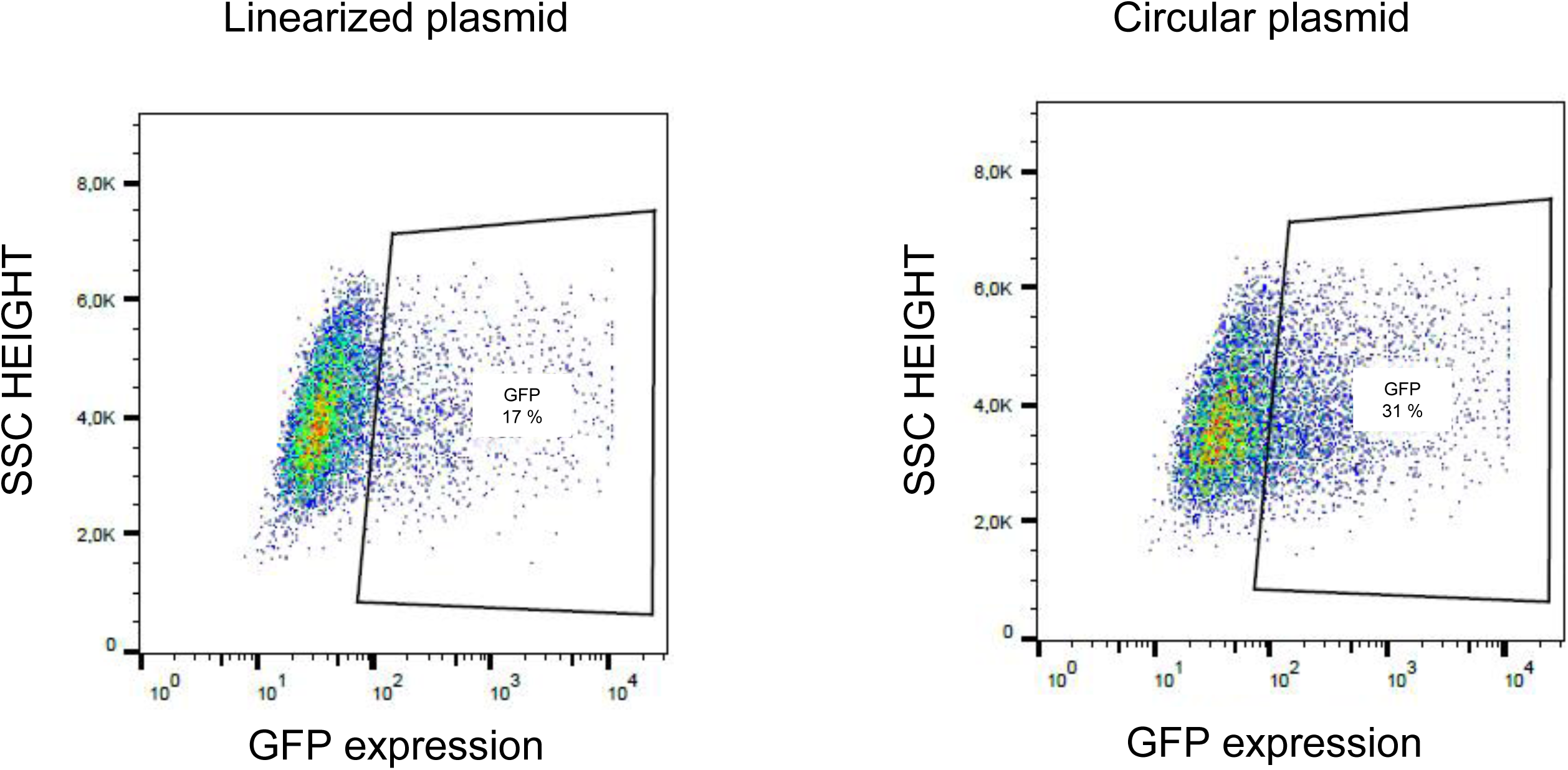
Assessment of transfection efficiency. Cells were transfected by nucleofection with either linear or circular plasmids and analyzed at the indicated timepoints in Figure 3E. After 72 hours, cells were assessed for transfection efficiency. Representative flow cytometry plots show GFP expression in cells transfected with either the linearized plasmid (left) or the circular plasmid (right). GFP-positive populations are gated, and the percentage is calculated based on the proportion of GFP-expressing cells within the total cell population.

**Supplementary Figure 4.**
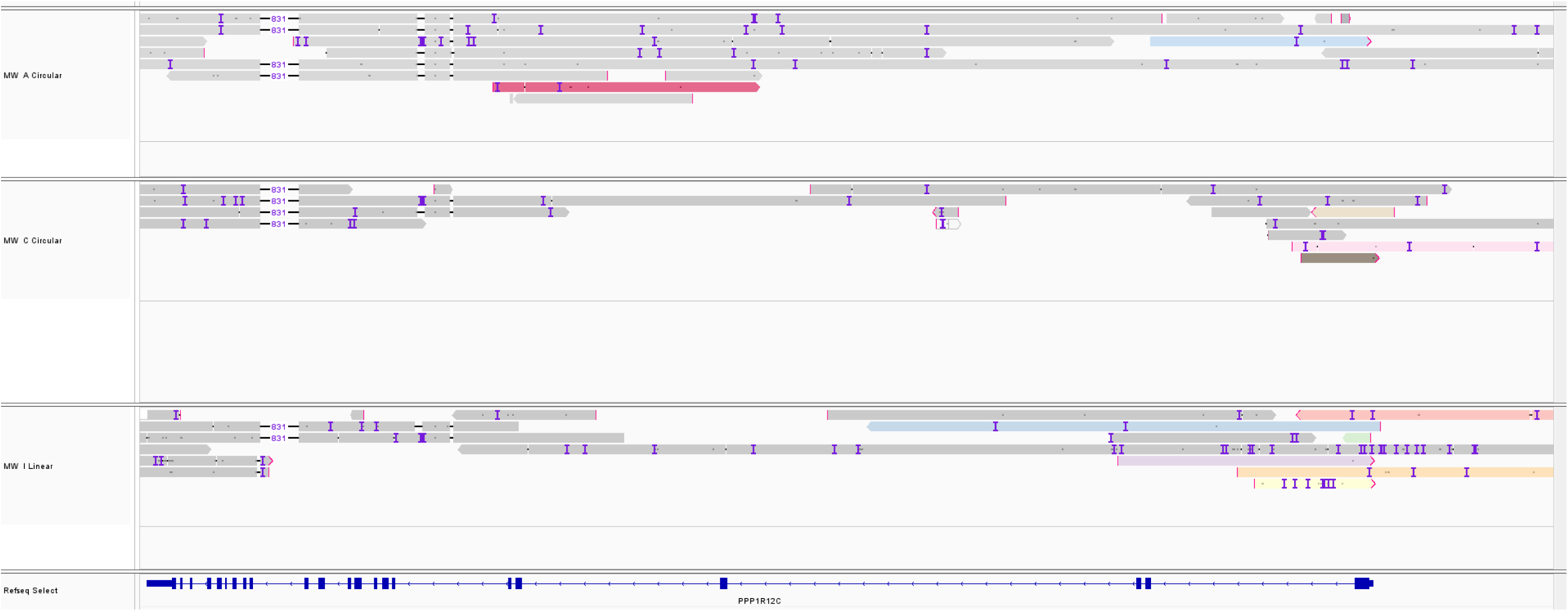
Nanopore sequencing alignment of selected MWs at the PPP1R12C locus (AAVS1) on chromosome 19. Long-read alignments from Oxford Nanopore sequencing are displayed for three MW: MW A (Circular), MW C (Circular), and MW H (Linear). Reads are aligned to the human reference genome (GRCh38), with the gene PPP1R12C annotated at the bottom (RefSeq track). Each row represents an individual read. Color-highlighted reads denote sequences that also align to the transgene insertion site, confirming breakpoint junctions or integration events. Small purple bars indicate mismatches, insertions, or deletions. The alignments span a ∼30 kb region within 19q13.42.

**Supplementary figure 5.**
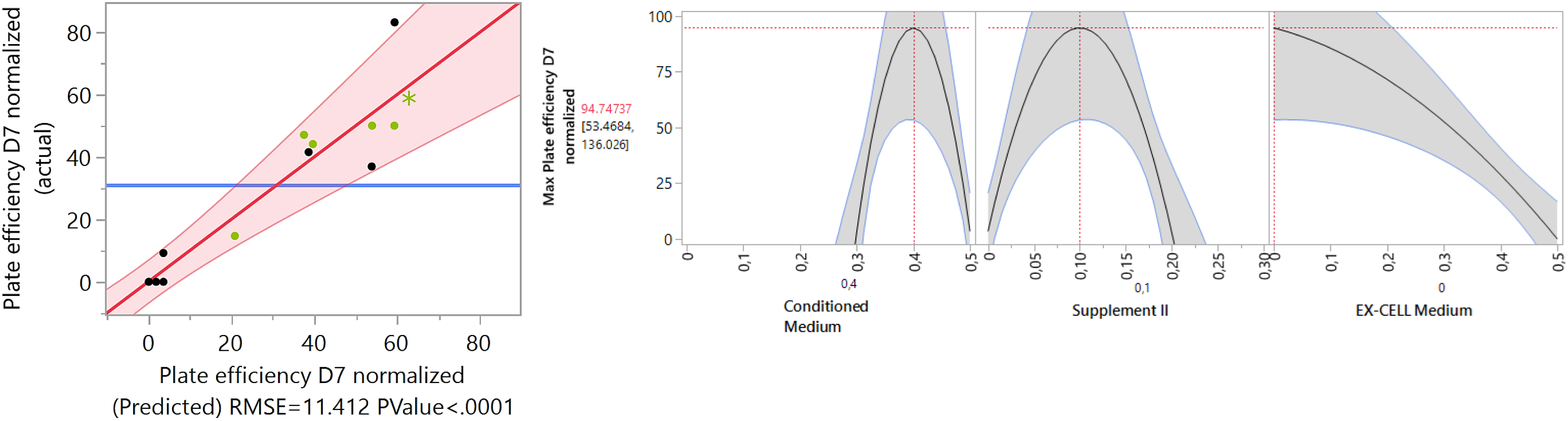
Optimization of media composition for colony formation. (A) Correlation between actual and predicted Percentage of wells with colonies at Day 7 (D7), based on a predictive model using media components as input variables. The shaded area represents the 95% confidence interval. The green points represent the plating efficiency on day 4. The model shows a good fit with RMSE = 11.412 and P-value < 0.0001. (B) Prediction profiler illustrating the effect of individual media components (Conditioned Medium, Supplement II, and EX-CELL medium) on plating efficiency. The profiler suggests an optimal combination with Conditioned Medium at ∼40%, Supplement II at ∼10%, and no EX-CELL medium.

